# Revealing COVID-19 Transmission by SARS-CoV-2 Genome Sequencing and Agent Based Modelling

**DOI:** 10.1101/2020.04.19.048751

**Authors:** Rebecca J Rockett, Alicia Arnott, Connie Lam, Rosemarie Sadsad, Verlaine Timms, Karen-Ann Gray, John-Sebastian Eden, Sheryl Chang, Mailie Gall, Jenny Draper, Eby Sim, Nathan L Bachmann, Ian Carter, Kerri Basile, Roy Byun, Matthew V O’Sullivan, Sharon C-A Chen, Susan Maddocks, Tania C. Sorrell, Dominic E Dwyer, Edward C Holmes, Jen Kok, Mikhail Prokopenko, Vitali Sintchenko

**Author notes:** these authors contributed equally (Rebecca J Rockett, Alicia Arnott). **Corresponding author**: Dr Rebecca Rockett, Sydney Medical School-Westmead, The University of Sydney, Australia, **Email**.

## Abstract

Community transmission of the new coronavirus SARS-CoV-2 is a major public health concern that remains difficult to assess. We present a genomic survey of SARS-CoV-2 from a during the first 10 weeks of COVID-19 activity in New South Wales, Australia. Transmission events were monitored prospectively during the critical period of implementation of national control measures. SARS-CoV-2 genomes were sequenced from 209 patients diagnosed with COVID-19 infection between January and March 2020. Only a quarter of cases appeared to be locally acquired and genomic-based estimates of local transmission rates were concordant with predictions from a computational agent-based model. This convergent assessment indicates that genome sequencing provides key information to inform public health action and has improved our understanding of the COVID-19 evolution from outbreak to epidemic.

---

In January 2020, a novel betacoronavirus (*Coronaviridae*), named Severe Acute Respiratory Syndrome coronavirus-2 (SARS-CoV-2), was identified as the etiologic agent of a cluster of pneumonia cases occurring in Wuhan City, Hubei Province, China, which were first reported in late December 2019^1,2^. The disease arising from SARS-CoV-2 infection, Coronavirus disease 2019 (COVID-19), subsequently spread rapidly worldwide. The World Health Organization (WHO) declared COVID-19 a pandemic on March 11^th^ 2020, when 118,000 cases had been reported from 110 countries. As of 18^th^ April 2020, the number of global cases had surpassed 2,000,000, following multiple worldwide independent importations of infection from visitors and returned travellers making the control of this disease of prime global public health importance^3,4^. Major outbreaks have been documented in South Korea, Iran, the USA and Europe^2,3^. At the time of writing, person-to-person transmission had been documented primarily through household contacts^5^, with up to 85% of human-to-human transmission occurring in family or household clusters^6,7^. The rapid growth in the number of COVID-19 cases with associated morbidity and mortality has overburdened healthcare facilities and the workforce. However, our understanding of the natural history and mechanisms of disease spread remain limited.

These events, combined with estimations from epidemic models, have led to unprecedented measures of disease control being instituted by national governments with profound costs to citizens and economies. Epidemic models of COVID-19 have suggested that virus transmission can be significantly disrupted by rapid detection and quarantine of infectious cases and their close contacts^8^. However, validation of COVID-19 modelling predictions is becoming increasingly important since many are built using incomplete and inconsistent data and thus produce divergent outcomes^9,10^, thus affecting confidence in public health policy directions. Here, we use the combination of near real-time SARS-CoV-2 genomic and public health surveillance data to verify inferences from computational models. Genomic epidemiology has become a high-resolution tool for public health surveillance and disease control^11-13^ and the COVID-19 pandemic has triggered unrivalled efforts for the real-time genome sequencing of SARS-CoV-2. Indeed, thousands of SARS-CoV-2 genomes have already been sequenced and made publicly available on GISAID (the Global Initiative on Sharing All Influenza Data)^14^. Importantly, the ongoing analysis of this global data set suggests no significant differences or links between SARS-CoV-2 genome sequence variability and virus transmissibility or disease severity^15^. However, even during these early stages of the global pandemic, genomic surveillance has been used to differentiate currently circulating strains into distinct, geographically based lineages and reveal multiple SARS-CoV-2 importations into geographical regions of China and the USA^16,17^.

Australia, as an island country between the Pacific and Indian oceans with strong traffic of people to and from COVID-19 hotspots in Asia, Europe and North America, has experienced unique challenges and opportunities in responding to the pandemic. The first laboratory-confirmed COVID-19 patients were diagnosed in Melbourne and Sydney on 25^th^ and 26^th^ of January 2020, respectively. Since then, and as of the end of this study period on March 29^th^ 2020, 4159 cases had been confirmed in Australia with 1981 cases (47.6%) occurring in New South Wales (NSW), the most populous state of Australia (24.5/100,000 population)^18^. The Australian Government introduced progressive epidemic mitigation measures on 23^rd^ March 2020 to limit social interactions, reduce virus diffusion and prevent community-based transmissions. This strategy has been supported by widely available testing for SARS-CoV-2 in NSW, with 1541 tests performed per 100,000 residents^19^.

In this study, we examine the value of near-real time genome sequencing of SARS-CoV-2 in understanding of local transmission pathways during the containment stage of the COVID-19 epidemic and compare findings from the genomic surveillance of SARS-CoV-2 with predictions of a computational agent-based model. This comparison was performed to assess the impact of potential sampling bias in genomic surveillance as well as to validate model-based inferences using experimental data. The synergistic use of high-resolution genomic surveillance and computational agent-based modelling not only improves our understanding of SARS-CoV-2 transmission chains in the community and the evolution of this novel virus but is essential for helping mitigate community-based transmissions.

## Results

### Sampling COVID-19 cases in the first phase of the Australian epidemic

Between January 26^th^ and March 28^th^, 1617 cases of COVID-19 were diagnosed and reported to the NSW Ministry of Health. All patients resided in metropolitan Sydney. Prior to February 29^th^, only four cases of COVID-19 were detected in NSW, all of which were imported. The first locally acquired case in NSW was reported on March 3^rd^, following which a sharp spike in both imported and locally acquired cases occurred during the week commencing March 15^th^ (Fig. 1). Between March 1^st^ and 21^st^, the weekly proportion of imported cases was between 5 and 20%. During the same period, cases epidemiologically defined as ‘unknown origin/under investigation’ increased from none during the week beginning March 1^st^ to between 31 and 35% from March 8^th^ to 21^st^ (Fig. 1C).

**Figure 1.**
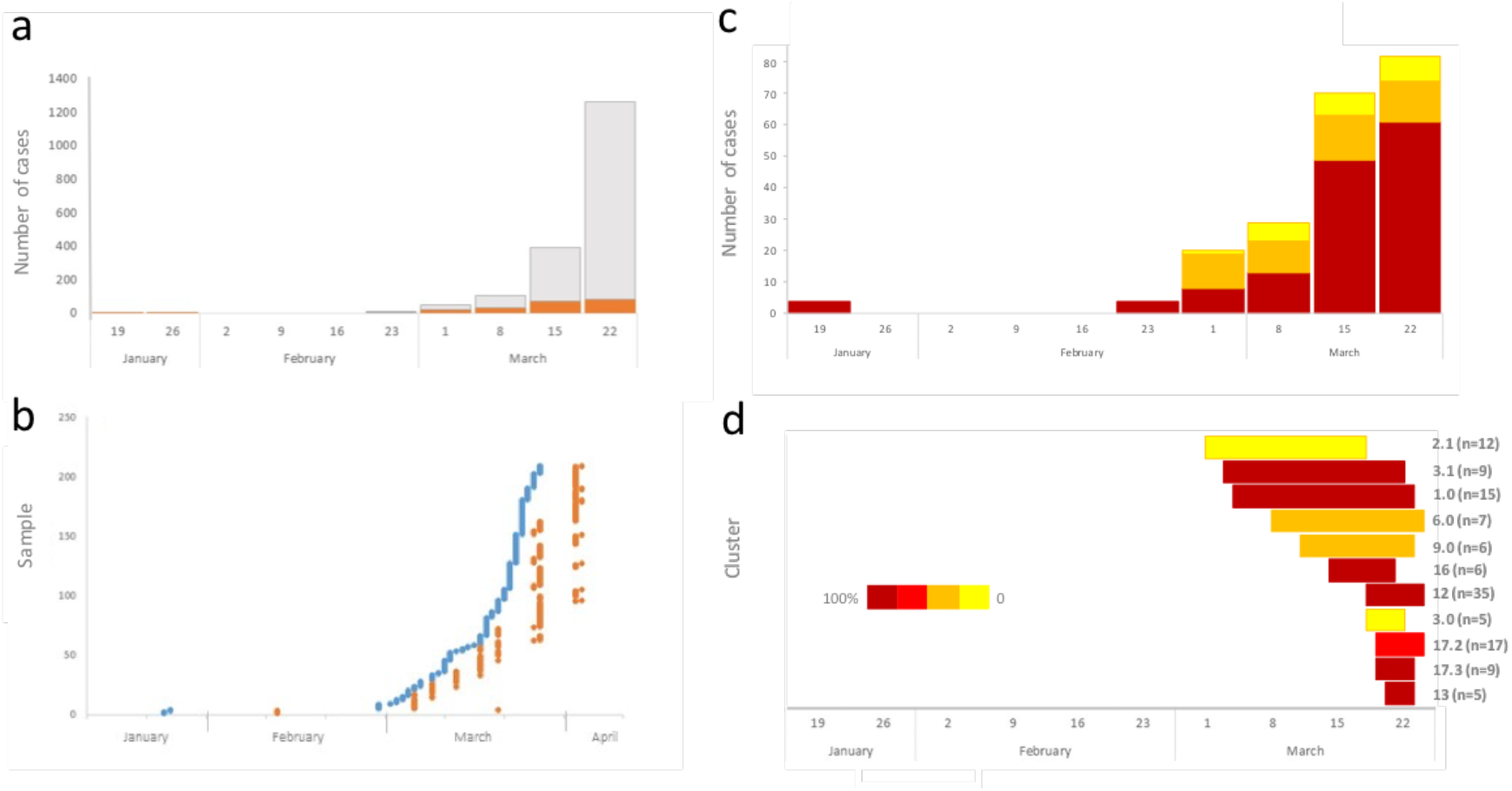
The timeline of COVID-19 cases and SARS-CoV-2 genome sequencing in NSW. (**a**) Weekly proportions of confirmed COVID-19 cases (grey) that underwent genome sequencing (orange) during the study period. (**b**) Turn-around-time of genome sequencing during the first phase of the epidemic. Shown are the date of collection (blue) and date of WGS (orange) for each of the 209 samples included in this study. (**c**) Total counts and proportions of imported (red) and locally acquired cases, both epi linked (orange) and of unknown source (yellow) during the study period by epi week. (**d**) Timelines of genomically defined clusters. Cluster number and number of cases (in brackets) are indicated next to bars (only clusters containing five or more cases are shown). Colour gradient reflects proportion of overseas acquired cases within the cluster, with darkest red representing 100% overseas acquired cases and yellow representing zero overseas acquired cases in the cluster.

### Rapid high-throughput SARS-CoV-2 sequencing directly from clinical samples

Of the 1,617 COVID-19 cases reported during the study period, complete viral genomes were obtained from 209 (13%) (Fig. 1a). Following an initial delay of 21 days between date of collection and the first sequencing run for the first three samples received, the median number of days between clinical sample collection and sequencing was five days (range: 1 – 21 days; Fig. 1b). The proportion of COVID-19 cases sequenced weekly peaked during the week beginning March 1^st^, at 25%. Consensus genome sequences were obtained from all specimens with RT-PCR Ct values <30. The amplicon sequencing method utilised in this study provided consistently complete, high quality consensus sequences with a median coverage depth between 277.4 and 3409.1 (Supplement Table S1). No significant changes in the amplicon primer sites were detected during the study.

### Evaluating phylogenetic diversity of SARS-CoV-2 genomes in NSW in January-March 2020

The 209 NSW SARS-CoV-2 genomes were dispersed across the global SARS-CoV-2 phylogeny (Fig. 2a). Notably, the majority of genomes from imported NSW cases grouped with viral sequences from the country of origin. All major genomic lineages of SARS-CoV-2 were detected in NSW (Fig. 2a, Supplement Table S1), including the introduction and then local transmission of lineage B.4.2 which is dominated by genomes from Australia that are believed to be imported from Iran (*24*).

**Figure 2.**
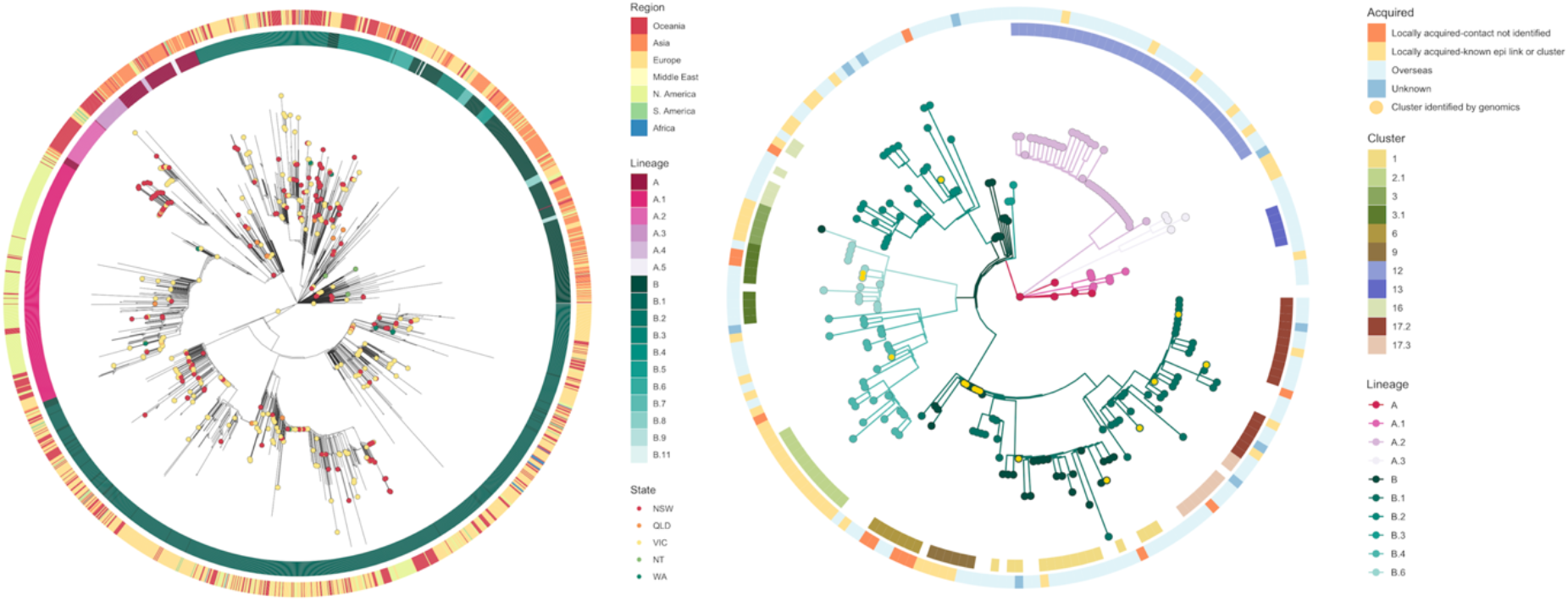
Phylogenetic analysis of SARS-CoV-2 genomes. (**a**) SARS-CoV-2 genomes from Australia in the global context provided by GISAID. Australian SARS-CoV-2 genomes are colour coded by the state of the initial diagnosis. The inner ring represents global phylogenetic lineages inferred from the final alignment using PANGOLIN (https://github.com/hCoV-2019/pangolin). Outer ring demonstrates the country of origin of GISAID genomes. (**b**) Phylogenetic relationships between SARS-CoV-2 genomes recovered from patients in NSW. The inner ring represents the allocation of clusters in NSW (only clusters equal or larger than 5 genomes are presented). Outer ring demonstrates the classification of cases as locally or overseas acquired based on genomic and epidemiological data. Bootstrap data for Lineage inference is presented in Supplemental Table S1.

### Close genomic similarity of SARS-CoV-2 in epidemiologically linked cases

Phylogenetic analysis identified multiple independent introductions of SARS-CoV-2 into NSW over time. SARS-CoV-2 from 10 patients (five household contacts pairs) were indistinguishable (three of five pairs) or differed by a single SNP (two of five pairs). However, institutional outbreaks, where cases could have only contracted the virus within the institution, demonstrated more genomic diversity. Three institutional outbreaks representing 35, 17 and 12 cases demonstrated up to two SNP differences between genomes during the outbreak. The duration of these three institutional outbreaks was between six and 17 days. We therefore chose to define clusters as sequences that differ by no more than 2 SNPs from the index case of each outbreak. In this study 27 clusters were identified, of which eight (29.6%) consisted of two cases and 11 (40.7%) consisted of five or more cases. All clusters consisted of five or more cases were associated with different institutions with no overlapping epidemiological connections. The largest cluster contained 35 cases linked by COVID-19 exposure in a single institution (Fig. 2b). With a single exception, all clusters remained active during the study period (Fig. 1d).

The phylodynamics of the epidemic in NSW was also investigated (Fig. 3a), however, genomic clusters were sampled for a limited period (maximum 19 days) and displayed weak temporal structure (R^2^=0.171). The tracking of cluster evolution over time will become increasingly important to identify active clusters over a longer sampling period. The low genetic diversity of SARS-CoV-2 in the early phase of this epidemic means that both genomic and epidemiological data are needed to clearly define SARS-CoV-2 outbreak clusters.

**Figure 3.**
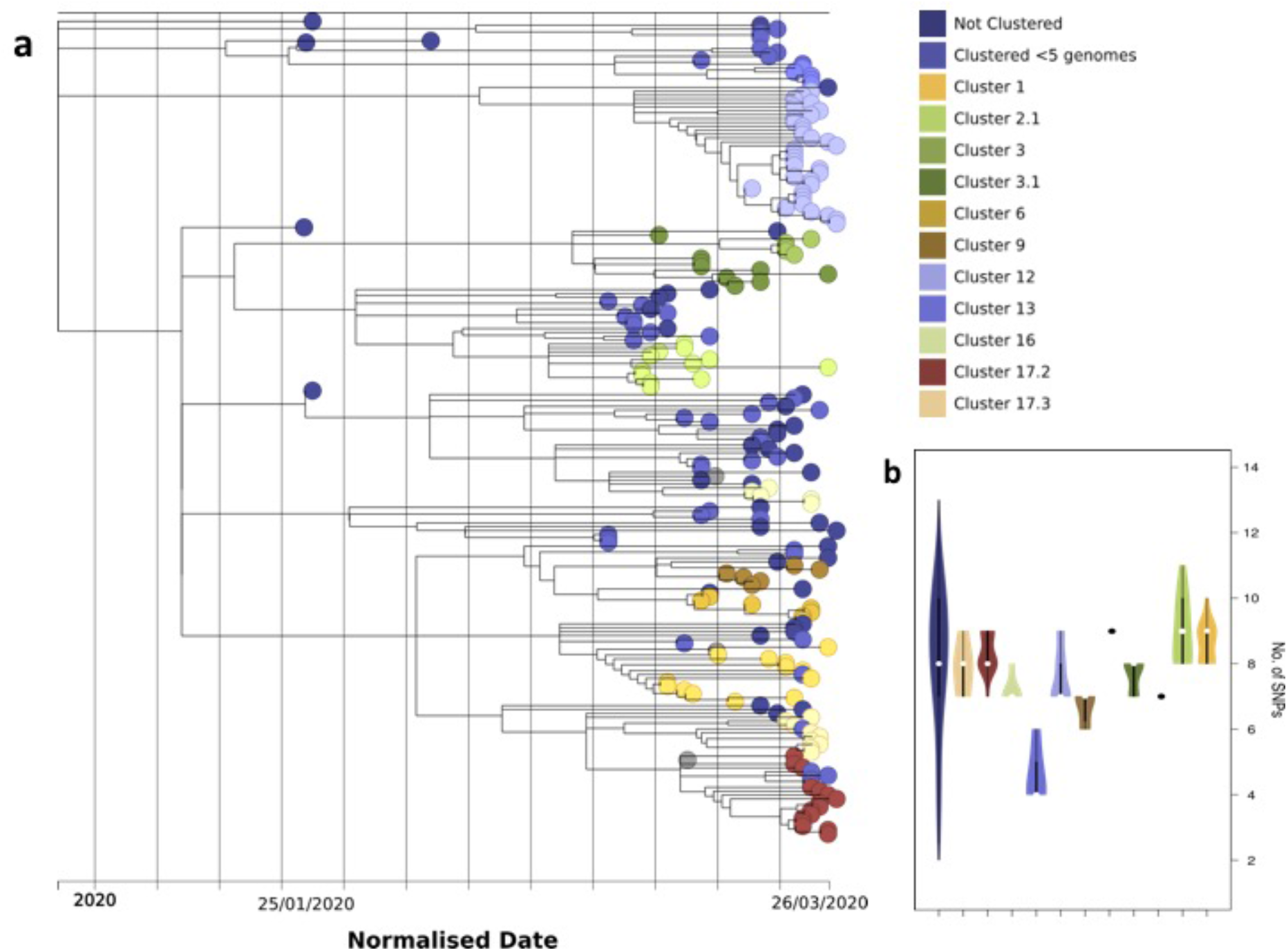
Phylodynamics of SARS-CoV-2 clusters in NSW. (**a**) Timestamped tree with selected genomic clusters (five or more cases); first Australian case was used as reference point; (**b**) Box plot contrasting SNP differences between selected genomics clusters and not clustered genomes of SARS-CoV-2 in NSW.

### Using genomics to cluster locally acquired cases with unknown epidemiological links

Twenty-two (10.5%) of the 209 cases included in this study were epidemiologically classified as ‘locally acquired – contact not identified’. Of these 22 cases, 15 (68%) were identified by genomic surveillance as belonging to nine genomic clusters containing cases with known epidemiological links (Fig. 2b). The remaining seven cases were found to be genomic singletons, not clustering with genomes included in this analysis.

### Agent-based model estimation of local transmission in Australia

In the agent-based model, the COVID-19 pandemic spread in Australia was initiated by overseas passenger arrivals, with some infections probabilistically generated in proportion to the average daily number of incoming passengers at airports, and binomially distributed within a 50km radius of each airport. Fig. 4a presents a network formed by community transmission chains produced by an ABM run simulating the period corresponding to the time interval between week 6 and week 10 of the study, that is, the period preceding the introduction of major lockdown strategies. A typical distribution of chain lengths is shown in Fig. 4.b, displaying similar trends in distribution of the genomic cluster sizes. The average local transmissions (within HH and HC contexts), obtained over ten simulation runs, amount to 18.6% (range 15.0-23.9%, with standard deviation 2.9%) of all infections simulated over the 35-day period, increasing over the weeks to 11.9% (range 8.0-14.7%, with std dev 1.9%) during the last week (week 10). When community transmissions are considered within a wider scope of local government areas, the average local transmissions (across HH, HC and SLA contexts) reach an upper bound of 34.9% (std dev 8.2%) of all infections, peaking during the last week at 24.4% (std dev 5.7%). These fractions are also found to be in strong concordance with their counterparts defined through the genomic cluster analysis (25.8% for all local transmissions, with 17.1–18.3% during the final week) (Fig. 4c). As expected, only a proportion of inferred transmission chains were detected by genome surveillance based on identified COVID-19 cases (Fig. 4b).

**Figure 4.**
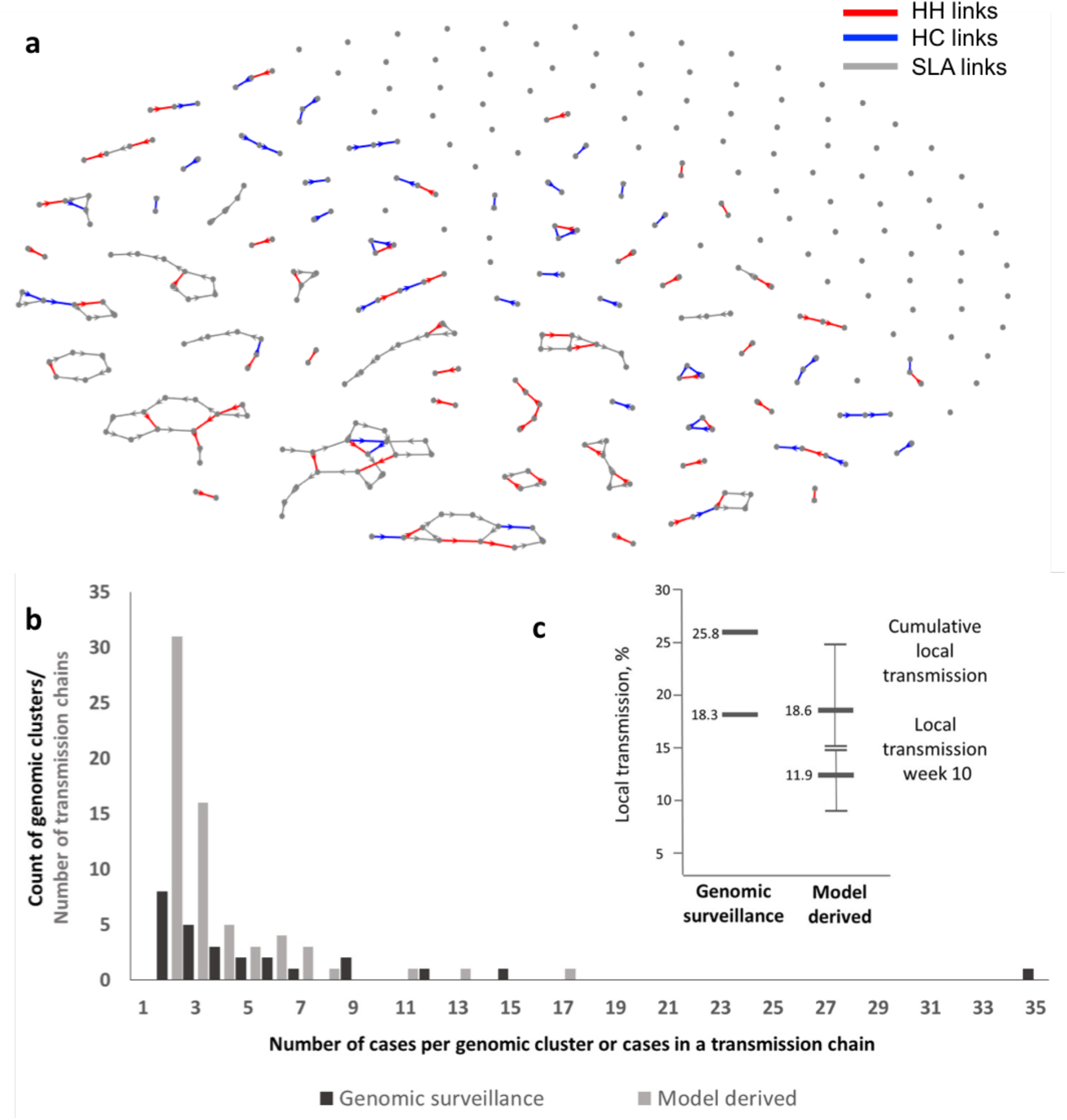
Network of SARS-CoV transmission cases generated by the agent-based model. (**a**) Network of transmission chains progressing through households (HH) and household clusters (HC) to statistical local areas (SLA). Local transmissions were defined as chains originated by these initial infections, and progressing through households and household clusters and mixing into wider neighbourhood, represented by statistical local area. (**b**) A typical distribution of chain lengths contrasted with sizes of genome sequencing defined clusters. (**c**) Comparison of local transmission rates in NSW defined by SARS-CoV-2 genome sequencing and simulated by the ABM model, averaged over ten simulation runs.

## Discussion

This is the first report of the convergent application of SARS-CoV-2 genomic surveillance and agent-based modelling to investigate the local transmission of COVD-19. Particular strengths of this study were integration of high-resolution genomic data with local epidemiology data and inferences made by agent-based modelling, providing context and confirmation for the genomic results and clustering. Our prospective SARS-CoV-2 genome sequencing has been instrumental in not only defining local transmission events and clusters, but enabling 68% of the cases for which no epidemiological links had been identified to be assigned to known epidemiological clusters, thereby allowing more efficient public health follow-up. The fine scale resolution provided by the genomic analyses presented in this study will become increasingly important for the containment of local outbreaks by enabling identification of secondary cases and providing context for cases of community transmission without clear epidemiological links^20^.

The high diversity of SARS-CoV-2 in Australia can be explained by multiple recent, concurrent and independent introductions of COVID-19 from countries with a high frequency of local transmission of COVID-19, such as Asia, Western Europe and North America. The decision to close the Australian border to all except Australian citizens, residents and immediate family members was announced on March 20^th^. Our phylogenetic analysis of unique sequences suggested or confirmed the country of origin for some cases and identified genomically clustered cases often associated with epidemiologically linked cases (Fig. 1b). This analysis documented genomically similar cases associated with concurrent community-based transmission in several institutions within the first ten weeks of the epidemic in Australia. However, it was not possible to infer a directionality of virus spread during these early stages of the epidemic as the rate of virus transmission is evidently greater than the rate of evolutionary change in the virus genome^16^. Our genomic data have also confirmed the dominance of overseas acquired infections in these early stages of the epidemic in Australia despite initial border control measures.

This observation based on the first ten weeks of the SARS-CoV-2 epidemic warrants further data collection in order to capture the impact of border control and social distancing measures introduced during the time of this study. The timing of this study, unfortunately, was also too short to delineate active (i.e. with continuing onward transmission) and inactive clusters. Although the genomic surveillance results are based on a relatively small number of COVID-19 cases in one Australian state, they offer a high-resolution picture of virus spread in the community and the concordance of results from genomic surveillance and agent modelling reinforces our conclusions.

This study leveraged the high resolution of inferences offered by agent-based infectious disease models in contrast to the simulation of the population-level dynamics models which have been typically employed for public health policy decisions. When such models are forced to rely on incomplete and inconsistent data, as is the case for the COVID-19 pandemic, they may suggest divergent outcomes, hence affecting public confidence in policy directions. The agent-based models capture interactions of individuals in time and space using government census data and detailed transmission pathways. The convergent estimates of local transmission from the computational model and genome sequencing not only validated COVID-19 modelling predictions but also suggested that our sampling of COVID-19 cases have been adequate for discovery and monitoring local transmission events in the time of cocirculation of genomically similar strains of SARS-CoV-2.

Our findings extend the value of genomic surveillance in depicting transmission pathways and evolution of emerging pathogens^13,21,22^. They also improve our understanding and support the implementation of genomically enhanced surveillance for more efficient COVID-19 control strategies and the investigation of cases with unclear infection sources and within a short turn-around-time^16,17,21,22^. The initial delay of 21 days between date of collection and sequencing for the first three samples received included the time taken to establish the in-house amplicon-based WGS method used in this study. The power of such surveillance is augmented by international sharing of SARS-CoV-2 genomic data collected by researchers and public health microbiology service providers^23,24^. Our clinical specimens originated from a mixture of private and public pathology providers, thus our sample likely represented a general population with a spectrum of clinical disease. We relied on an average sequence depth greater than 200 to ensure quality of short read mapping and genome comparisons in line with previous reports^20,25^, although further optimisation of may be required for sustainable and cost-effective SARS-CoV-2 surveillance.

In conclusion, this genomic survey of SARS-CoV-2 in a subpopulation of infected patients in the first ten weeks of COVID-19 activity in Australia enabled proactive monitoring of local transmission events in the critical time of implementation of national COVID-19 control measures. Only a quarter of cases appeared to be locally acquired and genomics-derived local transmission rates were concordant with predictions from the computational agent-based model. This convergent assessment improves our understanding of the COVID-19 evolution from outbreak to epidemic. Integrated analysis of outputs from SARS-CoV-2 genomic surveillance and computational models can refine our understanding of the evolution of COVID-19 epidemic and will be equally relevant for assessment of other emerging pathogens that public health is going to face in the future. Moving forward, in order to contain SARS CoV-2 in a relatively low-burden setting such as Australia, application of this high resolution genomic analysis will be crucial to track, trace and place cases in context to ensure targeted and informed public health action.

## Methods

### Samples for genomic surveillance

All clinical respiratory samples collected between January 21^st^ and March 25^th^, 2020, which tested positive by real time polymerase chain reaction (RT-PCR) for SARS-CoV-2 at the Institute of Clinical Pathology and Medical Research (ICPMR) were included in the study. The RT-PCR assay utilised WHO recommended primers and probes targeting regions in the E gene and RdRp domain (*20*). Samples were submitted directly to ICPMR for RT-PCR testing and also referred from other NSW Health Pathology and private pathology laboratories for genome sequencing. Genome sequencing was attempted on all PCR positive samples with a RT-PCR Ct value <30.

### Epidemiological data and case definitions

Public health follow-up was conducted in parallel for each COVID-19 case. Travel histories were collected and used to determine whether infections were acquired locally or overseas. The origin and number of confirmed COVID-19 cases detected in NSW during the study period were publicly available and obtained from the NSW Health website^26^.

An imported case was defined as a person who tested SARS-CoV-2 RT-PCR positive and reported international travel in the 14 days before the onset of illness. A locally acquired case was a person who tested SARS-CoV-2 RT-PCR positive and had not travelled outside of Australia in the 14 days before illness onset. Locally acquired cases were further classified on epidemiological grounds as confirmed contacts of a case, as part of a known cluster, or of unknown origin (i.e. no known contact with a case). Close contacts were defined as individuals who had been in the same closed space for at least two hours with a laboratory-confirmed case (i.e. someone who tested positive for COVID-19) when that person was infectious^27^.

The decision to close the Australian border to all except Australian citizens, residents and immediate family members was announced on March 20^th^. Stage 1 and 2 restrictions were implemented on March 23^rd^ and 26^th^ respectively. Stage 1 restrictions included closure of all restaurants, gymnasiums, cinemas and places of worship, as well as avoiding non-essential travel, outdoor gatherings of >500 and indoor gatherings of >100 people. Stage 2 restrictions were an extension of Stage 1 and included advising the public to stay at home unless going to work or education, shopping for essential supplies, undertaking personal exercise or attending medical appointments or compassionate visits^28^.

### SARS-CoV-2 genome sequencing

We undertook SARS-CoV-2 WGS using an existing amplicon-based Illumina sequencing approach^29,30^. Briefly, RT-PCR positive samples were reverse transcribed using SuperScript IV VILO MasterMix (ThermoFisher Scientific). The viral cDNA was used as input for multiple overlapping PCR reactions (~2.5kb each) that spanned the viral genome using Platinum SuperFi MasterMix. Amplicons were then pooled equally, purified and quantified before Nextera XT library preparation and multiplex sequencing on an Illumina iSeq or MiniSeq (150 cycle flow cell)^31^. All consensus SARS-CoV-2 genomes identified in the study have been uploaded to GISAID (Supplement Table S1).

### SARS-CoV-2 genome analysis

The raw sequence data was subjected to an in-house quality control procedure prior to further analysis. Demultiplexed reads were quality trimmed using Trimmomatic (sliding window of 4, minimum read quality score of 20, leading/trailing quality of 5)^32^. The taxonomic identification of the sample was verified using centrifuge version 1.0.4, based on a database compiled from human, prokaryotes and viral sequences (including SARs-CoV-2 Refseq sequences available prior to March 2020)^33^. All samples had > 99% assignment to genus ‘betacoronavirus’. Reference mapping and consensus calling was performed using iVar version 1.2^29^. Briefly, reads were mapped to the reference SARS-CoV-2 genome (NCBI GenBank accession MN908947) using BWA-mem version 0.7.17, with unmapped reads discarded. iVar trim was used to soft-clip reads containing primer sequences, and discard reads <20 length discarded following trimming. A consensus sequence was called for positions where depth >10, quality >20 with a minimum frequency threshold of 0.1. The 5’ and 3’ UTR regions were masked from the consensus due to poor quality of these regions. QUAST version 5.0.2 was used to evaluate the consensus sequence quality in addition to manual inspection in Geneious Prime (2020.0.5)^34^. SARS-CoV-2 genomes from NSW were compared with one another as well as with complete, or near complete, global genomes available at GISAID (www.gisaid.org: accessed 28^th^ March 2020, see Supplementary Table S2 for complete list of international genomes used in this study). The quality of GISAID genomes was evaluated using QUAST, with sequences retained only if they were >28,000-bp in length and contained <0.05% missing bases (n=1,985 reference genomes). The GISAID and NSW genomes were aligned with MAFFT v7.402 (FFT-NS-2, progressive method)^35^. Genomes were trimmed to remove 5’ and 3’ untranslated regions. Phylogenetic analysis was performed using the maximum likelihood approach (IQTree v1.6.7 (substitution model: GTR+F+R2) with 1,000 bootstrap replicates^36^. SARS-CoV-2 genomic lineages were inferred using Phylogenetic Assignment of Named Global Outbreak LINeages (PANGOLIN) (https://github.com/hCoV-2019/pangolin). Total SNP numbers between the index SARS-CoV-2 genome from NSW (GISAID Accession: NSW01/EPI-ISL-407893) and each genome in the study was calculated using SNP-sites (excluding ambiguities)^37^. Temporal structure and distribution of genomic clusters in NSW was visualised using Treetime^38^. Phylogenetic trees were constructed using R package ggtree^39^.

### Agent-based model

We used an agent-based model (ABM) developed to trace the ongoing COVID-19 pandemic in Australia^40^. This model has been specifically calibrated to reproduce several key characteristics of COVID-19 transmission in Australia, including its reproductive number (R0 = 2.27), the length of incubation and generation periods (5.0 and 6.4 days, respectively), age-dependent fractions of the symptomatic cases (0.669 and 0.134, for adults and children, respectively) and the probability of transmission from asymptomatic/pre-symptomatic agents (0.3 of symptomatic individuals). The ABM comprises approximately 24 million software agents, each with attributes of an anonymous individual, matching key demographic statistics from the 2016 Australian Census data as well as other data from the Australian Bureau of Statistics (ABS) and the Australian Curriculum, Assessment and Reporting Authority (ACARA). Contact and transmission rates were set to differ across distinct social contexts, such as households, household clusters, local neighbourhoods, schools, classrooms and workplaces. The ABM simulation processes and updates agents’ states over time, starting from initial infections, seeded in international airports around Australia, using demographics and mobility layers previously developed and applied to trace influenza pandemics^41-42^. Local transmissions were defined as links in the chains originated by all initial infections detected within a given context, i.e., progressing through households (HH) and household clusters (HC). In other words, transmissions occurring across the wider community are not counted as local transmissions, but are still detected in order to investigate the overall connectivity of emerging local “clusters”. Thus, each local transmission chain may include links from several mixing contexts: HH, HC and wider neighbourhood, represented by statistical local area (SLA) mapped to Statistical Area Level 2 (SA2) in the 2016 census (SLAs are Local Government Areas or part thereof). The COVID-19 model has been validated with the most recent disease surveillance data, being able to predict both the incidence and prevalence peaks in early to mid-April 2020^40^.

### Ethics statement

Clinical specimens were routinely processed at the Institute of Clinical Pathology and Medical Research (ICPMR) and deemed not research. A non-research determination for this project was granted by the Health Protection NSW as it was a designated communicable disease control activity.

## Supporting information

Bioinformatic metrics of consensus SARS-CoV-2 genome sequences

International and Australian SARS-CoV-2 genomes from GISAID used in this study

## Data availability

The original/raw SARS-CoV-2 genome sequencing data will be available in the National Center for Biotechnology Information GenBank by the time of publication. Consensus genome sequences for national and international genomes are available from the GISAID, (the Global Initiative on Sharing All Influenza Data; www.gisaid.org) (see Supplementary Tables S1 and S2). The ABM data sources have been detailed elsewhere^40-42^.

## Code availability

There are no unique pipelines or source code developed for this project.

## ACKNOWLEDGMENTS

The members of the ICPMR SARS-CoV-2 Study Group include Linda Donovan, Shanil Kumar, Tyna Tran, Hossinur Rahman, Danny Ko, Tharshini Sivaruban, Andrew Ginn, Qinning Wang and Keenan Pey. The members of the ABM team include Nathan Harding, Cameron Zachreson and Oliver Cliff. The authors acknowledge the Sydney Informatics Hub and the use of the University of Sydney’s high performance computing cluster, Artemis. The authors are grateful to NSW Pathology partner laboratories, ACT Pathology, Douglass Hanly Moir, Australian Clinical Laboratories and Laverty Pathology for referring samples for genomic surveillance. Expert advice and epidemiological information provided by the NSW public health surveillance units at the Health Protection, NSW Health are also gratefully acknowledged. The authors are indebted to all researchers and their organisations who have kindly shared SARS-CoV-2 genome data on GISAID.

## AUTHOR CONTRIBUTIONS

Study concept and design by VS, RR, AA, DED, JK, JSE, ECH & MP. Sample processing and testing by RR, CL, VT, KAG, ES, NB, & IC. Sequencing and analysis by RR, CL, RS, VT, KAM, JSE, MG, JD, KB, RB, VS & ECH. Agent-based modelling by SLC and MP. Study coordination by VS, SCAC, MVOS, SM, TS, DED & JK. VS, RR and AA wrote the first manuscript draft with editing from ECH, JSE, TS, DED, SAC & MP. The final manuscript was approved by all authors.

## Supplementary Material

**Table S1. Bioinformatic metrics of consensus SARS-CoV-2 genome sequences from NSW that have been uploaded to GISAID**

**Table S2. International and Australian SARS-CoV-2 genomes from GISAID used in this study**

